# Integrating episodic and spatial context signals in the hippocampus

**DOI:** 10.1101/2024.06.24.600445

**Authors:** Alexander Nitsch, Naomi de Haas, Mona M. Garvert, Christian F. Doeller

## Abstract

Episodic and spatial memory are the two key components of the mnemonic system in humans. Episodic memory enables us to remember events from the past whereas spatial memory enables us to form a map-like representation of the environment. Interestingly, these seemingly different functions rely on the same brain structure: the hippocampus. Yet, how the hippocampus supports both at the same time remains unclear. Here, we tested the hypotheses that the hippocampus supports these two systems either via a common coding mechanism or via a parallel processing mechanism. To this end, we combined functional magnetic resonance imaging (fMRI) with an episodic life-simulation task and a spatial virtual reality task to manipulate episodic and spatial context associations of objects. We then investigated fMRI adaptation effects between these objects as a result of shared contexts. We found that the fMRI signal in the anterior hippocampus scaled with the combined prediction of shared episodic and spatial contexts, in line with the idea of a common coding mechanism. We found no evidence for a parallel processing mechanism, as there were no differences between episodic and spatial effects. The common coding effect for episodic and spatial memory dovetails with the broader notion of domain-general hippocampal cognitive maps.

## Introduction

Memory is essential for everyday life as it allows us to store and retrieve experiences and knowledge to guide actions. Two key components of our mnemonic system are episodic and spatial memory. Episodic memory enables us to remember events and situations from our past, for example when, where and how we spent our last vacation (Davachi, 2006; Tulving, 2002). Spatial memory enables us to form a map-like representation of our environment that can be used for navigation, for example when walking home from the train station (Burgess et al., 2002; Epstein et al., 2017; Hartley et al., 2014; Tolman, 1948). Interestingly, both episodic and spatial memory rely on the same brain structure: the hippocampus (e.g. Burgess, 2014; Burgess et al., 2002; Buzsáki & Moser, 2013; Eichenbaum & Cohen, 2014; Kühn & Gallinat, 2014; O’Keefe & Dostrovsky, 1971; Scoville & Milner, 1957; Squire, 2009). However, it remains elusive how exactly the hippocampus supports both episodic and spatial memory at the same time and whether both rely on the same hippocampal coding principles.

Here, we propose that previous findings in the memory literature broadly offer two hypotheses to address this question: the hypothesis of a parallel processing mechanism and the hypothesis of a common coding mechanism, yet a thorough comparison between the two hypotheses is still lacking. The idea of a parallel processing mechanism is that neuronal populations processing episodic and spatial memory are fundamentally distinct and differently distributed across the hippocampus (Burgess et al., 2002; Ezzati et al., 2016; Hirshhorn et al., 2012; Kühn & Gallinat, 2014; Nadel et al., 2013; Persson et al., 2018; Poppenk et al., 2013; Spiers et al., 2001; Strange et al., 2014). Previous studies suggest functional differences between the two hippocampal hemispheres (Burgess et al., 2002; Ezzati et al., 2016; Spiers et al., 2001) and along the longitudinal axis (Hirshhorn et al., 2012; Nadel et al., 2013; Persson et al., 2018; Poppenk et al., 2013; Strange et al., 2014) of the hippocampus: More specifically, these studies suggest that episodic memory is rather supported by the left and / or anterior hippocampus while spatial memory is rather supported by the right and / or posterior hippocampus. This differential pattern is also in line with an extensive meta-analysis of human neuroimaging studies on episodic and spatial memory (Kühn & Gallinat, 2014). However, it is noteworthy that studies included in this meta-analysis investigated only either episodic or spatial memory, with systematic differences between the two study types (e.g. learning word lists for episodic memory vs. navigation in a virtual environment for spatial memory). In contrast, the idea of a common coding mechanism is that episodic and spatial memory are processed in the same way and thus by the same neuronal populations within the hippocampus (Behrens et al., 2018; Bellmund et al., 2018; Eichenbaum, 2014, 2017; Eichenbaum & Cohen, 2014; Epstein et al., 2017; O’Keefe & Nadel, 1978; Olsen et al., 2012; Schiller et al., 2015). One prominent view is that both episodic and spatial memory rely on hippocampal cognitive maps that encode relationships between states, e.g., between events in episodic memory or between places in spatial memory (Behrens et al., 2018; Bellmund et al., 2018; Eichenbaum & Cohen, 2014; Epstein et al., 2017; O’Keefe & Nadel, 1978; Olsen et al., 2012; Stachenfeld et al., 2017; Tolman, 1948). This view therefore highlights commonalities between episodic and spatial memory processes. Furthermore, previous studies demonstrated that the same hippocampal cells can code for specific points in time and in space – either as time cells or as place cells (Eichenbaum, 2014; Kraus et al., 2013; MacDonald et al., 2011). Correspondingly, recent human neuroimaging studies showed similar effects of temporal (i.e., episodic) and spatial distance representations in the hippocampus (Deuker et al., 2016; Kyle et al., 2015; Nielson et al., 2015).

To date, it remains elusive which of the two hypotheses more accurately reflects concurrent processing of episodic and spatial memory in the hippocampus, as conclusions based on previous studies are complicated by methodological challenges. For example, previous studies investigated only one mnemonic system at a time, used different learning material and analysis methods or investigated intertwined episodic and spatial effects during navigation (e.g. Deuker et al., 2016; Dimsdale-Zucker et al., 2018; Kühn & Gallinat, 2014).

Here, we aimed to address this gap by testing predictions of both a parallel processing and a common coding mechanism for episodic and spatial memory in the hippocampus in a single experiment with comparable episodic and spatial tasks and the same analysis method. To this end, we combined functional magnetic resonance imaging (fMRI) with an episodic life-simulation task and a spatial virtual reality task. We used these tasks to manipulate episodic and spatial context associations of objects, while holding episodic information constant across different spatial contexts and vice versa. We then used fMRI adaptation analyses (Barron et al., 2016; Desimone, 1996; Grill-Spector et al., 2006) to investigate the effect of these context manipulations on the neural representation of objects, assuming that shared contexts between them would result in adaptation effects in neuronal populations processing episodic and / or spatial memory. In line with the hypothesis of a common coding mechanism, we found an fMRI adaptation effect in the anterior hippocampus that scaled with the combined prediction of shared episodic and spatial contexts. We found no evidence for a parallel processing mechanism, as there were no differences between episodic and spatial effects.

## Results

### Participants form strong episodic and spatial object-context associations

To compare neural processing of episodic and spatial memory, 34 participants performed an episodic life-simulation task and a spatial virtual reality task (Fig. 1). The goal of both tasks was to manipulate episodic and spatial context associations of objects (Fig. 1b-c). In the episodic task, participants learned to associate objects with two episodic contexts, i.e., two stories in the life of a fictional character (Fig. 1f). Each story consisted of a sequence of six object-associated actions (e.g. the object bookshelf associated with the action “read a book”; see Supplementary Fig. 1 for an overview of all object-associated actions). In the spatial task, participants learned to associate objects with two spatial contexts, i.e., two neighborhoods of a virtual city (Fig. 1g). Participants had to deliver objects to a target store in the respective neighborhood (see Supplementary Fig. 2 for a layout of the virtual city). In both tasks, participants learned the object-context associations through feedback (Fig. 1f, g). Eight objects were associated with one of the two episodic contexts and one of the two spatial contexts (Fig. 1b; regular objects). Two episodic control objects appeared in both contexts of the episodic task and two spatial control objects appeared in both contexts of the spatial task (see fMRI results below for the purpose of including these control objects). The general structure of the two tasks was as similar as possible (e.g. in terms of the trial structure, identical isolated object presentations during trials, memory tests after task blocks). Furthermore, we held episodic information constant across spatial contexts (i.e., cover story of object delivery to stores in spatial task) and spatial information constant across episodic contexts (i.e., in the action-videos in the episodic task, the object was always placed at the same location behind a wall; see Methods for task details).

**Fig. 1.**
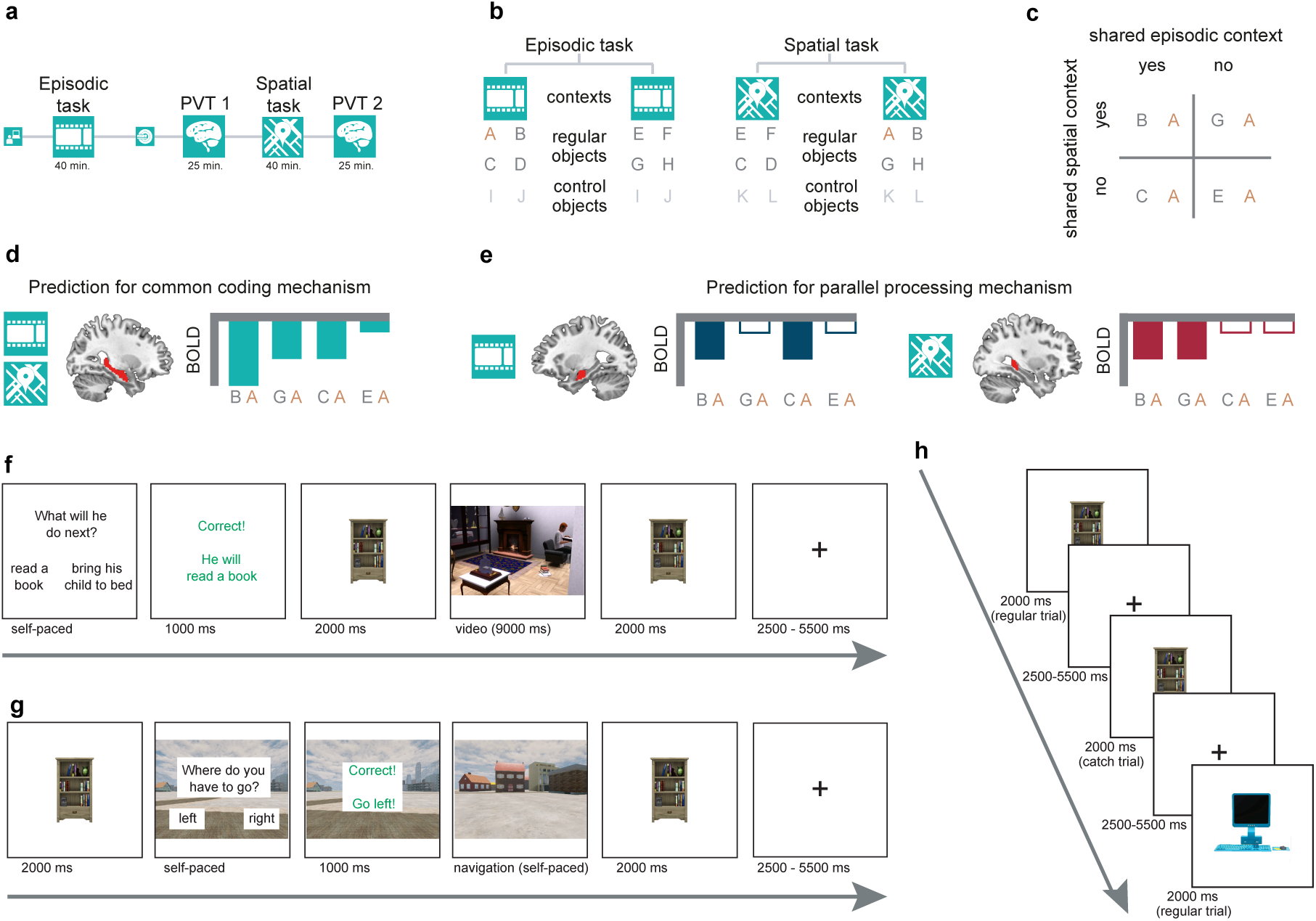
Study design and tasks. **a** Participants completed one of the two context association tasks, episodic or spatial, in a behavioral laboratory first and the other one in the scanner later (order counterbalanced over participants). Participants also completed two picture viewing tasks (PVTs) in the scanner, one after each context association task. **b** Both the episodic and the spatial task were divided into two contexts. Each regular object appeared in both tasks, but only in one of the two episodic contexts and in one of the two spatial contexts (e.g. object A). Control objects appeared in both contexts of a given task but did not appear in the other task (e.g. object I). **c** The distribution of objects across the task contexts resulted in a 2×2 design of pairs of objects sharing contexts (appearing in the same context). This included object pairs sharing both an episodic and a spatial context, object pairs sharing only one (either episodic or spatial) context and object pairs sharing no context. **d-e** To test predictions of a common coding and a parallel processing mechanism for episodic and spatial memory, we analyzed adaptation effects of these object pairs. For the common coding model (**d**), we predicted an adaptation effect scaling with the combined shared episodic and spatial contexts: highest adaptation effect for object pairs sharing both an episodic and a spatial context (e.g. BA), second highest for object pairs sharing only one – either a spatial context (e.g. GA) or an episodic context (e.g. CA) – and lowest for object pairs sharing no context (e.g. EA). For the parallel processing model (**e**), we predicted effects of shared episodic and spatial contexts to differ across subregions of the hippocampus (i.e., episodic effect stronger in the anterior and / or left hippocampus and spatial effect stronger in the posterior and / or right hippocampus). **f** Trial structure of the episodic task. In the episodic task, participants learned to associate objects with two stories (episodic contexts) in the life of a fictional character. Participants were asked to indicate the next object-associated action in a given story. They received feedback and saw the object. Afterwards, they watched a video of the fictional character performing the action before seeing the object again. **g** Trial structure of the spatial task. In the spatial task, participants learned to associate objects with two neighborhoods (spatial contexts) of a virtual city. Participants saw an object and were asked to indicate to which neighborhood they had to deliver the given object (sides of the neighborhoods were counterbalanced). After receiving feedback, they freely navigated to the target store in the neighborhood and finally saw the object again. **h** Trial structure of the PVT. Participants saw a stream of objects while performing a one-back cover task. They had to indicate whether the current object is the same as the preceding one (catch trials). **f-h** Stimuli of objects and videos in the episodic task were created using the computer game The Sims 3 (https://www.thesims3.com/). The virtual city was created using Unreal Development Kit 3 (Unreal Engine 3, Epic Games, Inc.).

We designed these tasks with the goal that participants form strong object-context associations, which was crucial for our planned fMRI analyses. Both tasks consisted of four learning blocks and participants completed a memory test after each block. In the memory test, they had to indicate for all object-context combinations of the respective task whether the object did or did not belong to the context (stories in the episodic task / neighborhoods in the spatial task). Participants remembered context associations of regular objects at ceiling level in the final memory tests of the episodic task (Fig. 2a; *M* = 99.45 %, *SD* = 2.33 %) and the spatial task (*M* = 98.90 %, *SD* = 3.55 %). There was no significant difference between the two scores (*t*(33) = 0.72, *p* = .66). Participants remembered context associations of control objects at ceiling level in the final memory test of the episodic task (*M* = 99.26 %, *SD* = 2.94 %) and very well in the spatial task (*M* = 89.71 %, *SD* = 19.29 %; significant difference between the two scores: *t*(33) = 2.90, *p* = .003). Across the four memory tests, participants reached ceiling performance in the episodic task earlier than in the spatial task (interaction task x block for regular objects: *F*(3,99) = 4.24, *p* = .007; interaction task x block for control objects: *F*(3,99) = 13.55, *p* < .001).

**Fig. 2.**
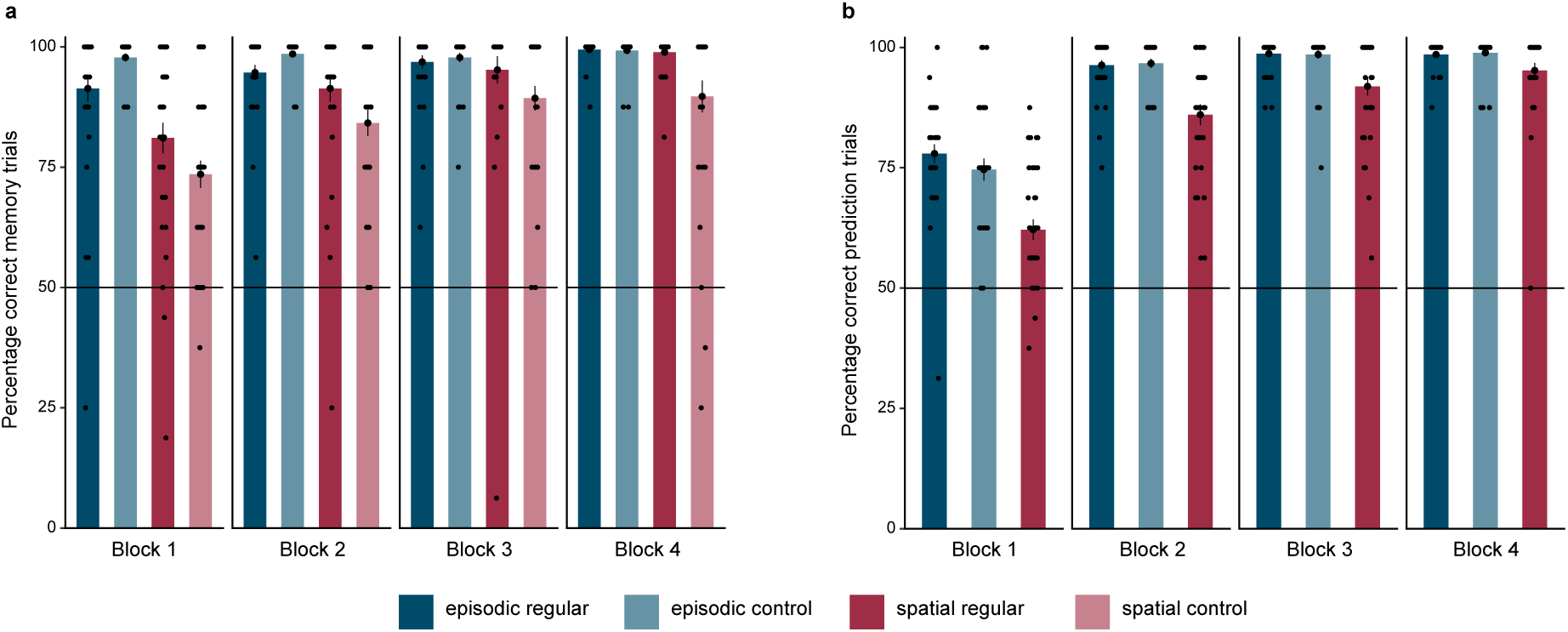
Participants form strong episodic and spatial context associations. **a** Performance in the memory tests after each task block. In the memory test, participants had to indicate for each object whether it belonged to a context. Performance is depicted separately for the episodic and spatial task and for regular and control objects. **b** Performance during prediction trials during each task block. In the episodic task, participants had to predict which object-associated action appeared next in the story. In the spatial task, participants had to predict to which neighborhood they had to deliver the current object. Performance is depicted separately for the episodic and spatial task and for regular and control objects (performance for spatial control objects is not depicted because both possible context (neighborhood) predictions were correct). **a,b** Dots represent data from *n* = 34 participants; bars and black circles with error bars correspond to mean ± SEM.

Participants’ strong memory is also evident during learning of the object-context associations, i.e., during the prediction trials of the episodic and spatial task. Participants performed at ceiling during the prediction trials in the final blocks of both the episodic task (Fig. 2b; regular objects: *M* = 98.53 %, *SD* = 3.05 %; control objects: *M* = 98.90 %, *SD* = 3.55 %) and the spatial task (regular objects: *M* = 95.22 %, *SD* = 9.23 %; control objects not scored because either answer was correct). Again, participants reached ceiling performance in the episodic task in earlier learning blocks than in the spatial task (interaction task x block for regular objects: *F*(3,99) = 7.76, *p* < .001).

Taken together, our behavioral results demonstrate that participants formed strong episodic and spatial object-context associations, enabling us to properly investigate and compare neural processing of episodic and spatial memory.

### Common coding and parallel processing mechanism for episodic and spatial memory in the hippocampus

To test our predictions of a common coding and a parallel processing mechanism using the same analysis method for both episodic and spatial memory, participants saw a stream of the objects in independent picture viewing tasks (PVTs), one after each context association task. To ensure that participants paid attention to the objects, they performed a one-back cover task, reporting whether the currently presented object is the same as the preceding one (Fig. 1h; performance PVT1: *M* = 86.92 %, *SD* = 17.88 %; performance PVT2: *M* = 91.50 %, *SD* = 14.26 %). The distribution of objects across the task contexts resulted in a 2×2 design of object pairs, with object pairs sharing both an episodic and a spatial context, object pairs sharing only one (either episodic or spatial) context and object pairs sharing no context (Fig. 1c). We investigated fMRI adaptation effects in the PVTs, with the idea that the fMRI signal in response to an object would be suppressed if there is high overlap between the neural representations of the current and the preceding object because of shared contexts (Barron et al., 2016; Desimone, 1996; Grill-Spector et al., 2006). This approach enables a clean and unconfounded comparison between episodic and spatial effects, as it is based purely on the presentation of the objects outside the task context and is therefore unaffected by differences in terms of visual complexity or any other potential remaining differences between the episodic and spatial task.

We first tested our prediction of a common coding mechanism that assumes that episodic and spatial memory are processed by the same neuronal populations. We thus hypothesized that, in regions with a common coding mechanism, the adaptation effect should scale with the combined prediction of shared episodic and spatial contexts after participants completed both association tasks (Fig. 1d). More specifically, the adaptation effect in PVT2 should be strongest for object pairs sharing both an episodic and a spatial context, lower for object pairs sharing only one – either an episodic or a spatial – context and lowest for object pairs sharing no context. We tested for such an adaptation effect by investigating whether the fMRI signal for an object was parametrically modulated by the number of episodic and spatial context associations between successively presented objects. In line with this common coding prediction, we observed a significant adaptation effect in the bilateral anterior hippocampus (Fig. 3a, b; small volume correction with *p_FWE_* < .05 TFCE; MNI peak voxel coordinates: -26,- 6,-20; peak voxel *t(*33) = -3.57, *p_FWE_* = .04; one-sided test). This effect cannot be explained by the number of context associations itself (no, one and two), since there was no significant effect in the cluster of the common coding effect for control objects which either share two context associations in the same task (episode or space) or no context association (Supplementary Fig. 4a; *t*(33) = -1.48, *p* = .08; one-sided test). Furthermore, the common coding effect for regular objects was significantly stronger than for control objects (*t*(33) = - 2.26, *p* = .03). These results show that the signal in the bilateral anterior hippocampus is consistent with a common coding mechanism for episodic and spatial memory.

**Fig. 3.**
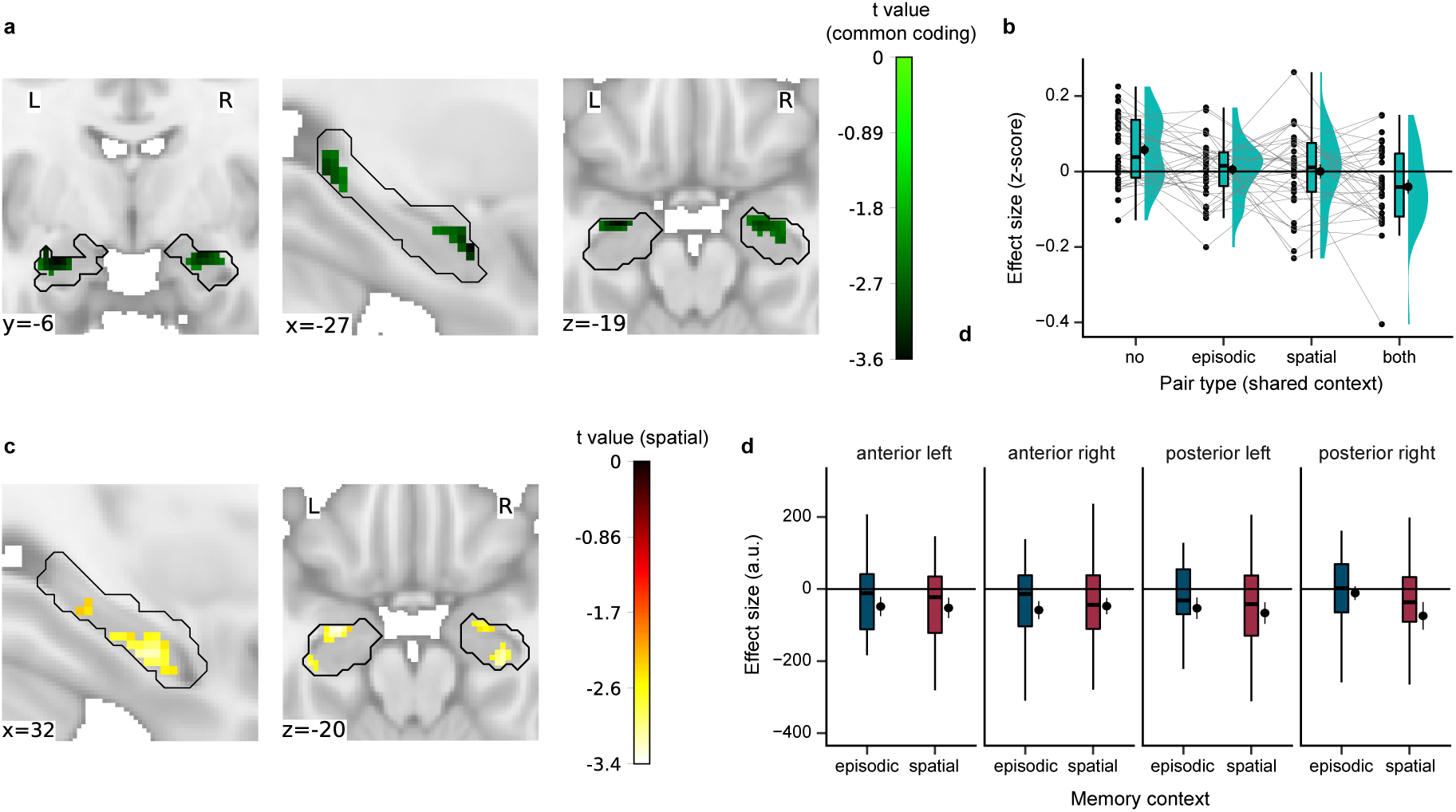
Common coding and parallel processing mechanism for episodic and spatial memory in the hippocampus. **a** Adaptation effect scaling with the combined prediction of shared episodic and spatial contexts, in line with a common coding mechanism for episodic and spatial memory. For illustration purposes, statistical image is thresholded at *puncorr* < .01; only the bilateral anterior hippocampus cluster survives correction for multiple comparisons (one-sided non-parametric permutation test with TFCE and small volume correction *pFWE* < .05). **b** Visualization of the adaptation effect in the common coding cluster from a. Depicted are effect sizes of the individual pair types, with the strongest adaptation effect for object pairs sharing both an episodic and a spatial context. **c** Spatial adaptation effect (spatial context shared vs. not shared). For illustration purposes, statistical image is thresholded at *puncorr* < .01; only a cluster in the right intermediate-anterior hippocampus survives correction for multiple comparisons (one-sided non-parametric permutation test with TFCE and small volume correction *pFWE* < .05). **d** Episodic and spatial adaptation effects, divided by hippocampal subregions with respect to the hemispheres and the longitudinal axis. There were no significant interactions between episodic and spatial effects with the hemisphere and longitudinal axis. **a-d** Negative values indicate an adaptation effect, i.e., suppressed response to the second object of a pair. **a,c** Black outline depicts the hippocampal mask used for small volume correction. Statistical images are displayed on the MNI template. **b,d** Dots represent individual participants’ data; boxplots show median and upper/lower quartile with whiskers extending to the most extreme data point within 1.5 interquartile ranges above/below the quartiles; black circles with error bars correspond to mean ± SEM; distributions depict probability density functions of data points.

Next, we tested our prediction of a parallel processing mechanism that assumes that neuronal populations processing episodic and spatial memory are differently distributed across hemispheres and/or along the longitudinal axis of the hippocampus (Fig. 1e). To test this prediction, we investigated whether the suppression of the fMRI signal for an object that shared vs. did not share a context with the preceding object differed depending on whether the shared context was episodic or spatial. We observed no significant difference between episodic and spatial adaptation effects (*p_FWE_* = .55). There was a significant spatial adaptation effect in the right intermediate-anterior hippocampus (Fig. 3c; small volume correction with *p_FWE_* < .05 TFCE; MNI peak voxel coordinates: 28,-22,-16; peak voxel *t(*33) = -3.23, *p_FWE_* = .04; one-sided test) but no significant episodic adaptation effect (*p_FWE_* = .10; one-sided test). In addition, we conducted a complementary ROI analysis with hippocampal subregions to test for differences between episodic and spatial effects with respect to the hemispheres and the longitudinal axis (four ROIs: left vs. right for the anterior and posterior hippocampus, respectively). We observed no significant interactions between episodic and spatial effects with the hemisphere or longitudinal axis (Fig. 3d; interaction memory x hemisphere x axis: *F*(1,33) = 1.96, *p* = .17; interaction memory x hemisphere: *F*(1,33) = 0.61, *p* = .44; interaction memory x axis: *F*(1,33) = 2.99, *p* = .09; interaction hemisphere x axis: *F*(1,33) = 1.71, *p* = .20; memory: *F*(1,33) = 0.43, *p* = .52; hemisphere: *F*(1,33) = 0.98, *p* = .33; axis: *F*(1,33) = 0.00, *p*

= .98).

Taken together, these results provide evidence for a common coding mechanism for episodic and spatial memory in the anterior hippocampus, with an adaptation effect scaling with the combined prediction of shared episodic and spatial contexts. We found no evidence for a parallel processing mechanism, as there was no difference between episodic and spatial effects.

### Episodic and spatial context signals on a whole-brain level

In this study, we primarily aimed to test our predictions of a common coding and a parallel processing mechanism for episodic and spatial memory in the hippocampus. Nevertheless, we also explored episodic and spatial effects on a whole-brain level. Interestingly, we observed an adaptation effect that scaled with the combined prediction of shared episodic and spatial contexts in a network of regions, most prominently in temporo-parietal regions (Fig. 4a; *p_FWE_* < .05 TFCE; one-sided test). We observed no significant difference between episodic and spatial effects, but separate significant episodic and spatial effects (Fig. 4b-c; *p_FWE_* < .05 TFCE; one-sided test).

**Fig. 4.**
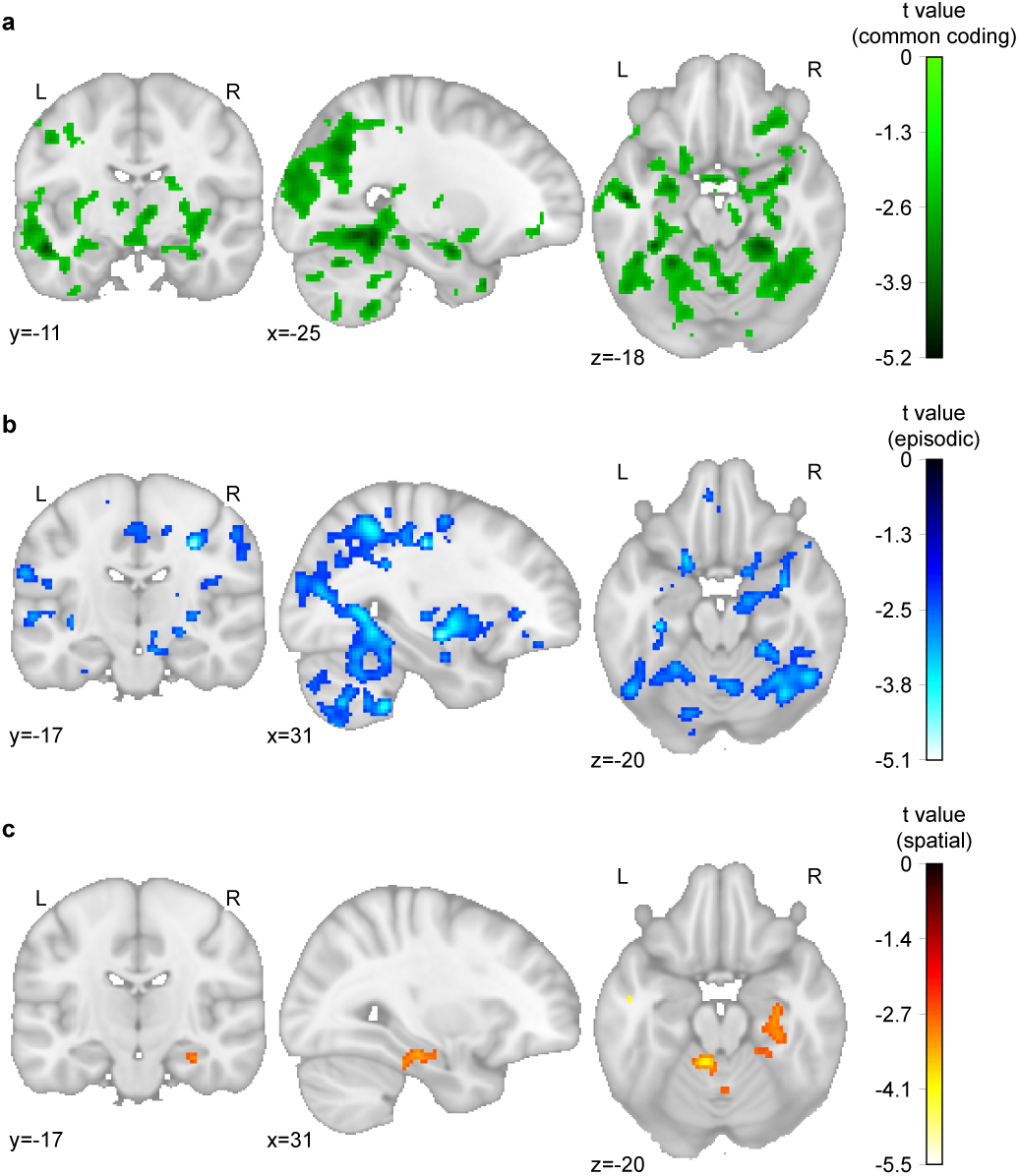
Episodic and spatial context signals on a whole-brain level. **a** Adaptation effect scaling with the combined prediction of shared episodic and spatial contexts (one-sided non-parametric permutation test with TFCE and *pFWE* < .05). **b** Episodic adaptation effect: episodic context shared vs. not shared. **c** Spatial adaptation effect: spatial context shared vs. not shared. **a-c** One-sided non-parametric permutation tests with TFCE and *pFWE* < .05. Negative values indicate an adaptation effect, i.e., suppressed response to the second object of a pair. Statistical images are displayed on the MNI template.

## Discussion

Episodic and spatial memory are core functions associated with the hippocampus. Yet how precisely the hippocampus can support both at the same time remains unclear. In this study, we set out to directly compare episodic and spatial memory effects in the hippocampus while testing the hypotheses of a common coding and a parallel processing mechanism. These hypotheses, both derived from the literature, assume that episodic and spatial memory are processed by either the same or by different neuronal populations within the hippocampus, respectively. Through comparable episodic and spatial tasks, participants learned to associate objects with episodic and spatial contexts, resulting in a 2×2 design of objects pairs sharing both an episodic and a spatial context, objects pairs sharing only one – either episodic or spatial – context and object pairs sharing no context. We then investigated fMRI adaptation effects between objects as a result of shared contexts. In line with the hypothesis of a common coding mechanism, we found an fMRI adaptation effect scaling with the combined prediction of shared episodic and spatial contexts in the anterior hippocampus. We found no evidence for a parallel processing mechanism, as there was no difference between episodic and spatial effects.

Our result of a common coding mechanism for episodic and spatial memory dovetails with the broader idea of hippocampal cognitive maps. Cognitive maps encode relationships between states in the world and are proposed to be domain-general (Behrens et al., 2018; Bellmund et al., 2018; Epstein et al., 2017; O’Keefe & Nadel, 1978; Schuck et al., 2016; Stachenfeld et al., 2017; Tolman, 1948; Wilson et al., 2014). In line with this, previous memory theories emphasized the relational structure of both episodic and spatial memory by binding together distinct elements across time and space into a common representation (Eichenbaum & Cohen, 2014; Olsen et al., 2012). These elements reflect events unfolding over time in episodic memory and places in space in spatial memory, both of which could be more generally understood as different instances of states in a cognitive map. Neurally, cognitive maps are assumed to rely on specialized cells in the hippocampal-entorhinal system (Behrens et al., 2018; Bellmund et al., 2018; Moser et al., 2017; O’Keefe & Nadel, 1978). For example, during spatial navigation, place cells exhibit increased firing at a particular location within an environment (O’Keefe & Dostrovsky, 1971). Time cells fire sequentially at specific time points of an experience (MacDonald et al., 2011). Interestingly, these cells also overlap, with the same cells coding for specific points in time and in space, as well conjunctively encoding time and spatial context as contextual time cells (Kolibius et al., 2023; Kraus et al., 2013; MacDonald et al., 2011; Omer et al., 2022). Furthermore, recent human neuroimaging studies show that hippocampal cognitive maps also represent abstract spaces and graph structures, underlining their role of a domain-general mechanism for representing task-relevant information (Bao et al., 2019; Constantinescu et al., 2016; Garvert et al., 2017, 2023; Nitsch et al., 2024; Tavares et al., 2015; Theves et al., 2019, 2020; Viganò et al., 2021, 2023).

The location of our common coding effect in the anterior hippocampus is also in line with previous studies showing temporal (i.e., episodic) and spatial distance coding in this region. For example, Deuker et al. (2016) let participants encounter objects while navigating along a route through a virtual city and found that pattern similarity of these objects in the right anterior hippocampus scaled with both temporal and spatial distances. Nielson et al. (2015) equipped participants with a lifelogging device and later showed participants pictures of their life events during fMRI. They found that pattern similarity in the left anterior hippocampus correlated with both temporal and spatial distances between these life events. Our results extend these reports of a spatiotemporal distance effect by showing that the anterior hippocampus also represents episodic and spatial context information – even without navigation in the episodic task. Furthermore, because we kept spatial information constant across episodic contexts (i.e., the object was placed at the same location in the videos) and episodic information constant across spatial contexts (i.e., cover story of object delivery to stores), our results reflect episodic context processing beyond the spatial domain and vice versa. In another study by Dimsdale-Zucker et al. (2018), participants viewed a series of videos (episodic contexts) showing first-person navigation through one of two houses (two spatial contexts). Objects were placed along the trajectory of the navigation in the video. Pattern similarity in left subfield CA1 was higher for object pairs that shared an episodic context than object pairs that did not share an episodic context. However, in this task design all objects that shared an episodic context also shared automatically a spatial context and the spatial context was not constant across all other episodes as in our design.

Our results provide no evidence for a parallel processing mechanism for episodic and spatial memory in the hippocampus, as there were no significant differences between episodic and spatial effects. The spatial effect in the right hippocampus itself is in line with previous studies relating spatial memory to the right hippocampus (Burgess et al., 2002; Ezzati et al., 2016; Kühn & Gallinat, 2014; Spiers et al., 2001). The idea of a parallel processing mechanism was supported in particular by a meta-analysis of human neuroimaging studies (Kühn & Gallinat, 2014). This meta-analysis found that episodic memory was rather supported by left and anterior hippocampal subregions and spatial memory was rather supported by right and posterior hippocampal subregions. However, as mentioned above, the studies included in the meta-analysis investigated only either episodic or spatial memory, which prevents a direct comparison of episodic and spatial memory that directly tests the two effects against each other. Furthermore, the two study types differed e.g. in terms of their learning material and analysis methods (e.g. learning word lists for episodic memory vs. navigation in a virtual environment for spatial memory). Here, we directly compared episodic and spatial effects using the same method of fMRI adaptation during object processing after the comparable episodic and spatial tasks in a single experiment. It is important to highlight that we examined adaptation effects in a separate picture viewing task, as this enabled a clean and unconfounded comparison. Thus, these effects most likely reflect automatic retrieval of the learned episodic and spatial contexts while viewing the objects. It is possible that the degree to which hippocampus processes episodic and spatial memory differently depends on current memory demands (e.g. whether it is an active task or whether a task encourages separation or integration of episodic and spatial information), in line with notions of the hippocampus flexibly shifting its representations depending on relevance (Abramson et al., 2023; Donoghue et al., 2023; Theves et al., 2020).

Lastly, our results dovetail with a body of literature showing that the hippocampus is crucial for context learning and context-dependent memory (Davachi, 2006; Hirsh, 1974; Julian & Doeller, 2020; Kennedy & Shapiro, 2004; Maurer & Nadel, 2021; Rugg et al., 2012). We speculate that episodic and spatial context learning in our study might also explain our exploratory whole-brain results, where a network of temporo-parietal to frontal regions showed an adaptation effect that scaled with shared episodic and spatial contexts. Many of these regions, e.g. precuneus and medial frontal gyrus, have been associated with context / source memory as well as with episodic and / or spatial memory in previous studies (Cavanna & Trimble, 2006; Duarte et al., 2010; Lie et al., 2006; Preston & Eichenbaum, 2013; Seger & Miller, 2010). Future studies could examine more precisely how these different regions interact during context processing. For example, a previous study showed that hippocampal context representations predicted the retrieval of associated task demands that were reinstated in dlPFC (Jiang et al., 2020). Taken together, our exploratory whole-brain results suggest that episodic and spatial context information is represented beyond the hippocampus in a network of temporo-parietal-frontal regions.

In conclusion, we set out to directly compare episodic and spatial memory in the hippocampus while testing the hypotheses of a common coding and a parallel processing mechanism in a single experiment. Our results are in line with the idea of a common coding mechanism in the anterior hippocampus, which dovetails with the broader idea of domain-general hippocampal cognitive maps.

## Methods

### Participants

38 participants were recruited online through the Radboud Research Participation System of Radboud University (Nijmegen, The Netherlands). Participants were screened through the recruiting system for fMRI exclusion criteria. Furthermore, experience with first-person computer games was recommended since one of the experimental tasks required navigation in a virtual city and might cause motion sickness. At the beginning of the study, all participants gave written informed consent and filled out an additional fMRI screening form.

Two participants stopped the experiment due to motion sickness after the spatial task and before the fMRI session. Two additional participants were excluded from the analyses, one due to mistakes during recording and one due to a memory score below 60% at the end of the spatial task. Thus, the final sample consisted of 34 participants (age: *M* = 23.21 years, *SD* = 3.36 years; 19 female; 18 participants with the episodic task first, 16 participants with the spatial task first).

At the end of the study, participants were reimbursed at a rate of 8 € / h for behavioral testing and 10 € / h for fMRI testing. The study took place at the Donders Institute - Centre for Cognitive Neuroimaging in Nijmegen (The Netherlands) and was approved by the local ethics committee (CMO Regio Arnhem-Nijmegen, The Netherlands, nb. 2014/288).

### Experimental procedure

#### Study design and general procedure

We combined fMRI with an episodic life-simulation task and a spatial virtual reality task to manipulate episodic and spatial context associations of objects. We then used fMRI adaptation analysis to investigate the effect of these context manipulations on the neural representation of these objects in independent picture viewing tasks (PVT).

More specifically, we manipulated context associations in a 2×2 design, whereby eight objects were associated with one of two episodic contexts (stories in the episodic task) and one of two spatial contexts (neighborhoods in the spatial task). This resulted in four different object pair types, with object pairs sharing both an episodic and a spatial context (appearing in the same episodic and the same spatial context), object pairs sharing either an episodic or a spatial context and object pairs sharing no context. We added four control objects: two episodic control objects appeared in both contexts of the episodic task and two spatial control objects appeared in both contexts of the spatial task. Episodic control objects did not appear in the spatial task and vice versa. Hence, control object pairs had the same number of context associations as object pairs sharing both an episodic and spatial context and therefore served as a control for an effect of association strength. Thus, there were twelve objects in total, with eight regular objects for the 2×2 design and four control objects. All objects were randomly assigned to the conditions for each participant. To create the object pictures, we used the computer game The Sims 3 (https://www.thesims3.com/) and Adobe Photoshop (https://www.photoshop.com/).

Our goal was to design the episodic and spatial task as comparable as possible. Thus, the general structure of the two association tasks was the same, i.e. the number of trials and blocks, duration, pseudo-randomization of ITIs, memory tests, and intermittent performance feedback (see below).

To investigate fMRI adaptation effects caused by the context associations, all objects were presented in pseudorandom order in independent picture viewing tasks (PVTs), one after each context association task.

The study lasted approximately 3 1⁄2 hours. Participants completed one of the context association tasks in a behavioral laboratory first and the other one then in the MRI scanner. The order of these tasks was counterbalanced across participants. The behavioral part took approximately 80 minutes. After a ten-minute break, participants continued with the MRI session for approximately 2 hours. In the scanner, participants performed three tasks: a first PVT (PVT1), the second context association task and a second PVT (PVT2).

#### Stimuli

Stimuli were pictures of everyday objects of the computer game The Sims 3 (https://www.thesims3.com/).

#### Episodic task

The goal of the episodic task was to associate objects with one of two episodic contexts. The eight regular objects were divided over the contexts so that four objects appeared in one context and the other four objects in the other context. The two episodic control objects were associated with both contexts. The two contexts were operationalized as two stories in the life of a fictional character. Each story consisted of a sequence of six object-associated actions (e.g. the action “read a book” associated with the object bookshelf). Participants were instructed to learn the sequence of the object-associated actions in each story. The sequence was fixed for a given participant but object-story associations were randomized across participants.

The task consisted of four blocks. In each block, each story was presented twice (two stories × six actions per story × two presentations = 24 trials per task block; 96 trials in the entire task).

A trial had the following structure:

1. At the beginning of each prediction trial, participants had to answer what the fictional character would do next (“What will he do next?”). Participants could choose between two possible actions by pressing one of two buttons on a keyboard or fMRI button box (self-paced). Participants had to guess the correct action at the beginning of the task and learned throughout the task. The given answer was highlighted for 0.5 s. The foil action was an action associated with one of the two stories. The side of the correct action was pseudorandomized so that for each action each side was equally often the correct answer. Furthermore, all actions appeared as answers as equally often as possible over the task, either as correct or foil answer. Secondly, for each correct action all other actions were equally often the foil answer as far as possible.
2. Feedback stating the correct action was presented for 1 s (e.g. “Correct! He will read a book” or “False! He will read a book”). Positive feedback was shown in green while negative feedback was shown in red.
3. The picture of the object associated with the action (e.g. bookshelf) was presented at the center of a white screen for 2 s.
4. A video showing the action was presented for 9 s (duration was chosen to approximately match the navigation time in the spatial task based on piloting data). Participants were instructed to consider the stories as plays with all actions taking place on the same stage. The physical layout (walls and floor) remained constant across all actions. However, the appearance of the stage changed for every action (e.g. living room or dining room). A thin strip on the right side of the stage was hidden by a wall. The action (e.g. reading a book) always took place on the same spot on the right side of the stage. The associated object (e.g. bookshelf) was always placed at the same location behind the wall in the right corner of the room. Therefore, the object was not visible to participants during the video. The rationale behind this cover story and this design was to hold any spatial information constant during the episodic task. This was important for the 2 × 2 design logic of object pairs in our study as the aim of the episodic task was to manipulate only episodic relations between objects. Videos were created using the computer game The Sims 3 (https://www.thesims3.com/).
5. The picture of the object associated with the action (e.g. bookshelf) was presented for a second time at the center of a white screen for 2 s.
6. A fixation cross appeared for a certain interstimulus interval (ITI). ITIs were jittered between one and three scanner pulses (TR was 1.5 seconds) plus 1 s (2.5 s, 4 s and 5.5 s). ITIs were pseudorandomized so that they appeared as equally often as possible for all actions. To avoid any imbalance in time between the four task blocks, the total length of ITIs within a task block had to stay within the range of one standard deviation of the mean of all total ITI lengths for the blocks.

The beginning of a story was signaled by the presentation of the name of the story for 1.5 s (“Story 1” or “Story 2”). After each story, feedback was presented for 5 s. The feedback stated the percentage of correct answers given within that story and a short motivation (“You scored […] % in the last block. Keep going!”).

After each task block, participants performed a short memory test (hence, four memory tests across the entire task). In the memory test, participants had to indicate for all objects for each story whether the given object did or did not appear in the given story (12 objects × two stories = 24 trials). Participants were informed that there might be objects that did not appear in either of the two stories (these were the spatial control objects). The given answer was highlighted for 0.5 s. At the end of the memory test, subjects received feedback for 3 s. The feedback stated the percentage of correct answers (“You scored […] % in the last block”) and a short motivation: “Perfect score!” in case of 100 %, “Great job! Try to get a perfect score next time” in case of more than 75 % and “Stay motivated and you can score even higher next time” in case of less than 75 %.

On average the whole task took 40.0 minutes (*SD* = 1.9 minutes). The task was programmed in Presentation 18.3 developed by Neurobehavioral Systems (https://www.neurobs.com/).

#### Spatial task

The goal of the spatial task was to associate objects with one of two spatial contexts. The eight regular objects were divided over the contexts so that four objects belonged to one context and the other four objects to the other context. The two spatial control objects were associated with both contexts. The two contexts were operationalized as two distinct neighborhoods in a virtual city. Participants were instructed to deliver objects to these neighborhoods.

The two neighborhoods, one mostly with skyscrapers and the other one mostly with one- and two-story houses, were separated by fallow land. On the fallow land, two identical warehouses were situated opposite each other. In each neighborhood, there was a store as the target location for the delivery. The stores looked identical. Furthermore, Euclidean distances between the warehouses and the stores were identical. Participants had to pick up objects from one of the two warehouses and deliver it to the target store of the corresponding neighborhood. The purpose of including two warehouses was to prevent pure object-side associations. The cover story of the delivery ensured that any episodic information was constant across the whole task. This was important for the 2 × 2 design logic of object pairs in our study as the aim of the spatial task was to manipulate only spatial relations between objects. Object-neighborhood associations were randomized across participants.

The task consisted of four blocks. Each block consisted of four delivery blocks with six trials (four delivery blocks × six trials = 24 trials per task block; 96 trials in the entire task). Regular objects had to be delivered equally often in each task block (two trials for a regular object in a task block). Furthermore, spatial control objects had to be delivered equally often to each neighborhood within a task block (two trials for one neighborhood and two trials for the other in a task block). Participants were instructed that for objects which were sold in both neighborhoods, each neighborhood was the correct delivery target for half of the trials of the given object in a task block.

A trial had the following structure:

1. The picture of the current object was presented at the center of a white screen for 2 s.
2. Participants were placed at one of the two warehouses and had to indicate whether the object had to be delivered to the neighborhood that was on the left or the right side of the current warehouse (question “Where do you have to go?”, with the scenery of the fallow land and the two neighborhoods in the background). Participants could rotate but not change their location. They indicated their response by pressing one of two buttons on a keyboard or fMRI button box (self-paced). Participants had to guess the correct neighborhood at the beginning of the task and learned throughout the task. The given answer was highlighted for 0.5 s. Both warehouses appeared equally often as starting location in each block and for every object-neighborhood association. The order of object deliveries was pseudorandomized under the aforementioned conditions.
3. Feedback stating the correct direction was presented for 1 s (e.g. “Correct! Go left” or “False! Go left”). Positive feedback was shown in green while negative feedback was shown in red.
4. Participants could freely navigate to the target store in the correct neighborhood by using the arrow buttons on a keyboard or a fMRI button box (approx. 10 s). Participants received warnings in case they navigated away instead of towards the correct target store. A critical distance to trigger a warning was defined as the Euclidean distance between the warehouse and the correct target store plus a third of this distance. Two possible scenarios could trigger warnings: participant walking towards the wrong target store (warning displayed in red with correct direction, e.g. “Go left!”) or by participant surpassing the target store and walking too far into the correct neighborhood (warning displayed in red: “Too far. Go back.”)
5. After walking into the target store the object counted as delivered and the picture of the object was presented for a second time at the center of a white screen for 2 s.
6. A fixation cross appeared for a certain interstimulus interval (ITI). ITIs were jittered between one and three scanner pulses (TR was 1.5 seconds) plus 1 s (2.5 s, 4 s and 5.5 s). ITIs were pseudorandomized so that they appeared as equally often as possible for all objects. To avoid any imbalance in time between the four task blocks, the total length of ITIs within a task block had to stay within the range of one standard deviation of the mean of all total ITI lengths for the blocks.

After a delivery block of six trials, feedback was presented for 5 s. The feedback stated the percentage of correct answers given within that delivery block and a short motivation (“You scored […] % in the last block. Keep going!”).

After each task block, participants performed a short memory test (hence, four memory tests across the entire task). In the memory test, participants had to indicate for all objects for each neighborhood whether the given object did or did not belong to the given neighborhood (12 objects × two neighborhoods = 24 trials). Participants were informed that there might be objects that did not belong to either of the two neighborhoods (these were the episodic control objects). The given answer was highlighted for 0.5 s. At the end of the memory test, subjects received feedback for 3 s. The feedback stated the percentage of correct answers (“You scored […] % in the last block”) and a short motivation: “Perfect score!” in case of 100 %, “Great job! Try to get a perfect score next time” in case of more than 75 % and “Stay motivated and you can score even higher next time” in case of less than 75 %.

Before the beginning of the task, participants completed a training of 3 minutes to familiarize themselves with the virtual city. In the training, participants could navigate freely in the virtual city and had to look for the warehouses and the stores. These were marked by a cone in front of them during training only. Participants who performed the spatial task in the scanner completed the training in the behavioral laboratory before.

On average the spatial task took 40.0 minutes (*SD* = 3.6 min). The task was programmed in Unreal Development Kit 3 (Unreal Engine 3, Epic Games, Inc.).

#### Picture viewing task (PVT)

The goal of the PVT was to independently investigate the effect of the episodic and spatial context associations on the neural representation of the objects using fMRI adaptation analysis. Participants completed two identical PVTs in the MRI scanner, one after each context association task.

Participants were instructed that they would see a stream of objects. To ensure that they paid attention to the objects, they performed a one-back cover task, comparing the current object to the preceding one. Participants had to press one of two buttons on a fMRI button box if the current object was the same as the preceding one (catch trials) and the other button if the objects were not the same. Button contingencies were randomized across participants. This one-back task was orthogonal to later analyses of the PVTs.

The twelve objects from the episodic and spatial tasks were presented during the PVTs at the center of a white screen. Each PVT consisted of 208 trials with a trial duration of 2 s. The order of the objects was pseudorandomized across participants (see below). However, the object order in PVT 1 and PVT 2 was identical for a given participant. Each PVT was divided into four blocks of 52 trials. After each block, participants had a 20 second break. For the first 15 s, they received feedback on the percentage of correct answers and a repetition of the instructions. Afterwards, they saw a countdown for 5 s before the next block. 24 of the 208 trials in the whole task were catch trials (self-repetitions, around 11.5% of all trials) and these trials were evenly distributed across the twelve different objects (two per object) and the four blocks (six per block).

The order of the objects was pseudorandomized with the purpose to maximize power for the planned adaptation analysis. The idea was that the fMRI signal in response to an object would be suppressed if there is high overlap between the neural representations of the current and the preceding object because of shared contexts (Barron et al., 2016; Desimone, 1996; Grill-Spector et al., 2006). Hence, we ensured that all objects were preceded by all other objects they formed a relevant pair with. Each block started with trials for regular objects (first 33 non-catch trials) and ended with trials for control objects (last 13 non-catch trials). This allowed us to have the maximum number of relevant object transitions, as we were not interested in adaptation effects between regular and control objects. Regarding the regular objects, there were four types of object pairs, with object pairs sharing both an episodic and a spatial context, object pairs sharing only an episodic context, object pairs sharing only a spatial context and object pairs sharing no context. The object order was pseudorandomized so that all types of object pairs were presented equally often in each block (eight times per type). Furthermore, all regular objects appeared four times in each block. For each pair type, each possible combination of objects was presented as equally often as possible within a block and across the whole task (maximum differences in combinations within type in each block was 1). Lastly, for each object pair, either object was the preceding one in two out of the four blocks (AB vs. BA). Regarding the control objects, there were three types of object pairs, with object pairs sharing two episodic contexts, object pairs sharing two spatial contexts and object pairs sharing no context. Each type of control pairs was presented four times in a block. Each control object appeared three times in a block. The rest of the pseudorandomization was analogous to the pseudorandomization of the regular objects.

After every object presentation, a black fixation cross was presented at the center of the screen for a certain ITI. ITIs were jittered between one and three scanner pulses (TR was 1.5 seconds) plus 1 s (2.5 s, 4 s and 5.5 s). ITIs were pseudorandomized so that they appeared as equally often as possible across all non-catch trials of an object (the ITI of a catch trial was randomly chosen). To avoid any imbalance in time between the four blocks, the total length of ITIs within a block had to stay within the range of one standard deviation of the mean of all total ITI lengths for the blocks.

The task was programmed in neurobs Presentation (version 16.4, www.neurobs.com/presentation).

#### MRI data acquisition

MRI data were recorded using a 3 Tesla Siemens Magnetom Prisma Fit scanner (Siemens, Erlangen, Germany) with a 32-channel head coil. Functional T2*-weighted images for the two PVTs and the second context association task were acquired with a 4D multiband sequence with 84 slices (multi-slice mode, interleaved), TR = 1500 ms, TE = 28 ms, flip angle = 65 deg, acceleration factor PE = 2, FOV = 210 mm and an isotropic voxel size of 2 mm. A T1-weighted MPRAGE anatomical image was acquired with TR = 2300 ms, TE = 3.03 ms, flip angle = 8 deg, FOV = 256 x 256 x 192 mm and an isotropic voxel size of 1 mm. If the time limit of 2 hours for the scanning session was not yet reached, two separate phase and magnitude images were acquired in order to correct for distortions with a gradient field map (multiband sequence with TR = 1020 ms, TE1 = 10 ms, TE2 = 12.46 ms, flip angle = 45 deg, voxel size of 3.5 x 3.5 x 2.0 mm).

### Behavioral data analysis

We performed all behavioral analyses using MATLAB R2019b (https://de.mathworks.com/products/matlab.html) and Python 3.8 using Spyder (https://www.spyder-ide.org/; version 5.1.5) distributed via Anaconda (https://www.anaconda.com/; version 2020.11). Statistical analyses were based on the Python packages scipy (version 1.10.0) and statsmodels (version 0.13.2). T-tests and correlations tests were based on non-parametric permutation-based approaches to assess significance (10000 permutations). If not stated otherwise, we used an alpha level of .05 and two-sided tests.

We assessed performance as the proportion of correct trials and reaction times (log-transformed) in the memory and prediction trials of the episodic and spatial task – separately for the two tasks, regular vs. control objects and task blocks. We tested for performance differences in the final memory tests of the episodic and spatial task using related-samples *t*-tests. To assess learning, we also tested for performance and reaction time differences by block and task using a repeated measures ANOVA. Furthermore, we tested for performance differences between the four contexts (two episodic and two spatial contexts) in the final memory tests of the episodic and spatial task using a repeated measures ANOVA.

### fMRI data analysis

We performed all fMRI analyses using FSL (version 6.00, http://fsl.fmrib.ox.ac.uk/fsl/fslwiki/), MRIcron (Beta 2007, https://www.nitrc.org/projects/mricron), MATLAB R2019b (https://de.mathworks.com/products/matlab.html) and Python 3.8 in Spyder (https://www.spyder-ide.org/; version 5.1.5) distributed via Anaconda (https://www.anaconda.com/; version 2020.11). Statistical analyses were based on FSL Randomise as well as the Python packages scipy (version 1.10.0) and statsmodels (version 0.13.2). T-tests and correlations tests were based on non-parametric permutation-based approaches to assess significance (10000 permutations). If not stated otherwise, we used an alpha level of .05 and two-sided tests.

### Region of interest (ROI) definition

For our hippocampal small volume analysis, we used a bilateral hippocampal mask provided by the WFU pickatlas (Maldjian et al., 2003). Additionally, we used masks of four hippocampal subregions: anterior left, anterior right, posterior left and posterior right hippocampus. Following Collin et al. (2015) and Theves et al. (2019), the posterior portion ranged from Y = −40 to −30 and the anterior portion ranged from Y = −18 to −4 to increase sensitivity for differences between the anterior and posterior hippocampus.

### Preprocessing

We converted DICOM files of the MRI scanner to NIfTI files using MRIcron (Beta 2007, https://www.nitrc.org/projects/mricron). Subsequently, we preprocessed the data using FSL. We removed non-brain tissue from the structural images and applied motion correction and a high pass filter (threshold of 100 s) to the functional images. Furthermore, we coregistered structural and functional images using 6 DOF and a field map image if acquired for the participant. We registered structural images to the MNI152 template using 12 DOF and nonlinear registration. Finally, we registered the functional images to the participant’s anatomical space for the first-level analysis. Volumes that exceeded a movement cut-off of 4 mm and/or had artifacts (e.g. distortions throughout the whole brain) were modeled in later first-level GLMs with a volume-specific regressor, in addition to the six movement parameters.

### Analysis of episodic and spatial context signals

To test our predictions of a common coding and a parallel processing mechanism for episodic and spatial memory in the hippocampus using the same analysis method, we investigated fMRI adaptation effects between objects in the PVTs. The distribution of objects across the task contexts resulted in a 2×2 design of object pairs, with object pairs sharing both an episodic and a spatial context, object pairs sharing only one (either episodic or spatial) context and object pairs sharing no context. We thus investigated fMRI adaptation effects, with the idea that the fMRI signal in response to an object would be suppressed if there is high overlap between the neural representations of the current and the preceding object because of shared contexts (Barron et al., 2016; Desimone, 1996; Grill-Spector et al., 2006). In our first-level GLMs of the PVTs, we weighted transitions between object presentations according to the different models.

The GLMs contained separate onset regressors for each of the twelve objects to account for any object-specific differences in activity. Catch trials (self-repetition of an object) were modeled in separate regressors per object. In the GLM testing the common coding prediction in PVT2, each onset regressor was accompanied by a parametric regressor reflecting the weight given to the pair of the current and the preceding object according to the common coding model. The common coding model assumes that episodic and spatial memory are processed by the same neuronal populations. We thus hypothesized that, in regions with a common coding mechanism, the adaptation effect should scale with the combined prediction of shared episodic and spatial contexts after participants completed both association tasks. More specifically, the adaptation effect in PVT2 should be strongest for object pairs sharing both an episodic and a spatial context (weight: 4), lower for object pairs sharing only one – either an episodic or a spatial – context (weight: 3) and lowest for objects pairs sharing no context (weight: 2). Furthermore, we used control objects to test for a general effect of association strength. Control objects either share two context associations in the same task (episode or space) or no context association. The weights for control object pairs corresponded to the respective weights of regular object pairs (two shared: 4, not shared: 2). All parametric regressors were demeaned. To measure the adaptation effect of regular objects, the contrast weights of all parametric regressors of regular objects were set to 1. To measure the adaptation effect of control objects, the contrast weights of all parametric regressors of control objects were set to 1.

In the GLM testing the parallel processing prediction in PVT2, each onset regressor was accompanied by two parametric regressors reflecting the weight given to the pair of the current and the preceding object according to the shared episodic context (i.e., episodic context shared (weight: 2) vs. not shared (weight: 1)) and according to the shared spatial context (i.e., spatial context shared (weight: 2) vs. not shared (weight: 1)), respectively. All parametric regressors were demeaned. Contrast weights of the episodic parametric regressors were set to 1 to measure the episodic effect and contrast weights of the spatial parametric regressors were set to 1 to measure the spatial effect. To test for a difference between the two, contrast weights of the episodic parametric regressors were set to 1 and those of the spatial parametric regressors to -1. In the GLM testing the parallel processing prediction in PVT1 after participants completed only one context association task, each onset regressor was accompanied by a parametric regressor reflecting the weight given to the pair of the current and the preceding object according to the participant’s first context association task.

While these GLMs had the advantage that they accounted for any object-specific differences in activity, they lost information about the specific effects of the different pair types. We therefore used additional GLMs to visualize pair-specific effects. These GLMs contained separate regressors for each possible pair of regular objects and each possible pair of control objects. To further remove object-specific differences in activity from the resulting pair-specific parameter estimates, we demeaned each parameter estimate by the mean of the parameter estimates of the current object (second object in pair) and z-standardized them. To measure the effect of a pair type, we averaged the z-scores belonging to the given pair type.

All GLMs included regressors for the following events of no interest: two regressors for the two possible button presses with a stick duration, one regressor for the beginning of the task until the first object presentation, one regressor for the end of the task from the end of the last object presentation until end of scanning, one regressor for all break blocks.

The GLMs were computed in participants’ native space. We then spatially normalized the relevant outputs (contrast estimates and/or parameter estimates) to MNI standard space and afterwards smoothed them using a 6 mm full-width at half maximum Gaussian kernel for the group level analysis.

On the group level, we tested the significance of contrasts across participants using non-parametric permutation testing implemented in FSL Randomise with 10000 permutations. We used threshold-free cluster enhancement and corrected for multiple comparisons with family-wise error rate (*p_FWE_* < 0.05). We conducted analyses with small volume correction based on our a priori ROI of the hippocampus (see ROI definition) and additionally explored whole-brain effects. We used one-sided tests based on our a priori hypothesis of adaptation effects due to shared contexts between objects.

For the common coding effect in the anterior hippocampus, we controlled for a general effect of association strength by testing for an adaptation effect of control objects using a one-sample *t*-test (one-sided) and by testing for a difference between the common coding effect of regular objects and the adaptation effect of control objects using a related-samples *t*-test. For comparability of the two effects, we averaged the respective parameter estimates of the parametric regressors of the effects. Furthermore, we tested whether the common coding effect differed by group of participants having completed the episodic or spatial task first using an independent samples *t*-test. We also calculated a Spearman correlation between the common coding effect and performance in the episodic and spatial memory tests, averaged across tasks and blocks (note that performance was at ceiling in the final test).

For the spatial effect in the right hippocampus, we tested whether the effect differed by group and whether it was correlated with spatial memory test performance.

To test the parallel processing prediction, we also conducted a complementary ROI analysis with hippocampal subregions to test for differences between episodic and spatial effects with respect to the hemispheres (left vs. right) and the longitudinal axis (anterior vs. posterior). For this purpose, we extracted mean contrast estimates of the episodic and spatial effects of each ROI and then used a 2×2×2 repeated measures ANOVA. For both PVT 1 and PVT 2 we tested for the within-subject factors hemisphere and axis. For PVT 1, we added the between-subject factor group (episodic vs. spatial). For PVT 2, we added the within-subject factor shared context (episodic vs. spatial).

### Analysis of encoding-related activity during the context association tasks

To investigate encoding-related activity during the episodic and spatial context association tasks, the GLMs contained object-specific regressors for the different trial phases. For both the episodic and the spatial task, there was one regressor per object modeling the question and feedback and one regressor per object modeling the object presentation. In addition, the GLM for the episodic task contained one regressor per object modeling the video of the object-associated action and the GLM for the spatial task contained one regressor per object modeling the navigation to deliver the object. For both tasks, the GLMs contained regressors for button presses, for the feedback after every six trials and for the memory tests. To measure encoding-related activity, the contrast weights of the video regressors in the episodic task and the navigation regressors in the spatial task for regular objects were set to 1.

The GLMs were computed in participants’ native space, the contrast estimates spatially normalized to MNI standard space and smoothed using a 6 mm full-width at half maximum Gaussian kernel. On the group level, we tested the significance of contrasts across participants using non-parametric permutation testing implemented in FSL Randomise with 10000 permutations. We used threshold-free cluster enhancement and corrected for multiple comparisons with family-wise error rate (*p_FWE_* < 0.05).

Furthermore, we correlated encoding-related activity in the episodic and spatial tasks with the common coding effect in PVT2 in the anterior hippocampus (Pearson correlation). Note that participants performed only one context association task in the fMRI scanner so that we used the encoding effect of that task for a given participant (16 participants for the episodic task and 18 participants for the spatial task). For comparability of episodic and spatial encoding effects, we used t-values. Similarly, we correlated encoding-related activity in the spatial task with the spatial effect in PVT2 in the right hippocampus (Pearson correlation).

## Supplementary figures

**Supplementary Fig. 1.**
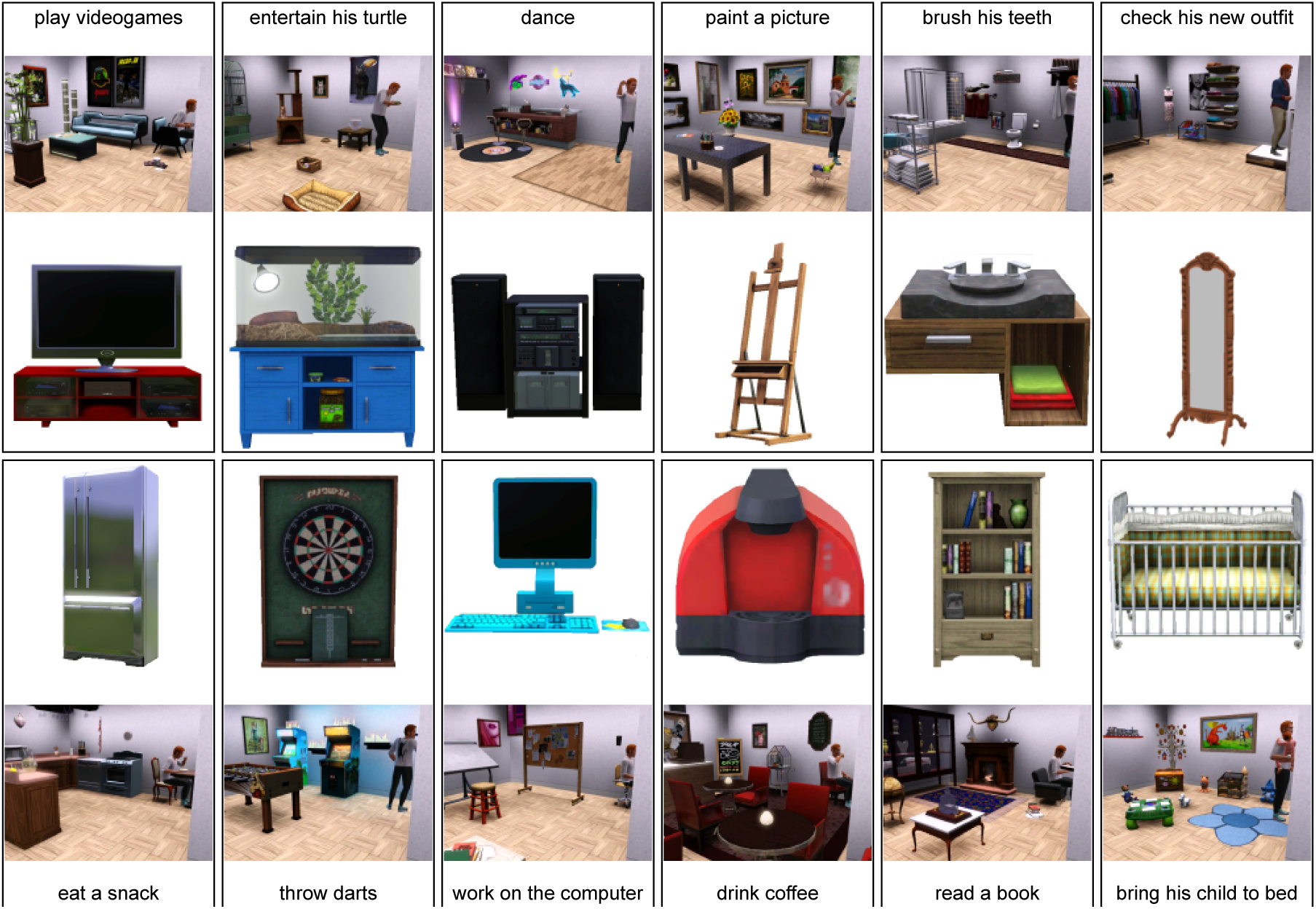
Objects and associated actions in the episodic task. Depicted are all twelve objects used throughout the experiment. In the episodic task, each object was associated with a corresponding action, depicted here by screenshots of these action-videos. Note that only ten of the twelve objects appeared in the episodic task for a given participant and the two remaining objects were spatial control objects. Stimuli of objects and the videos were created using the computer game The Sims 3 (https://www.thesims3.com/).

**Supplementary Fig. 2.**
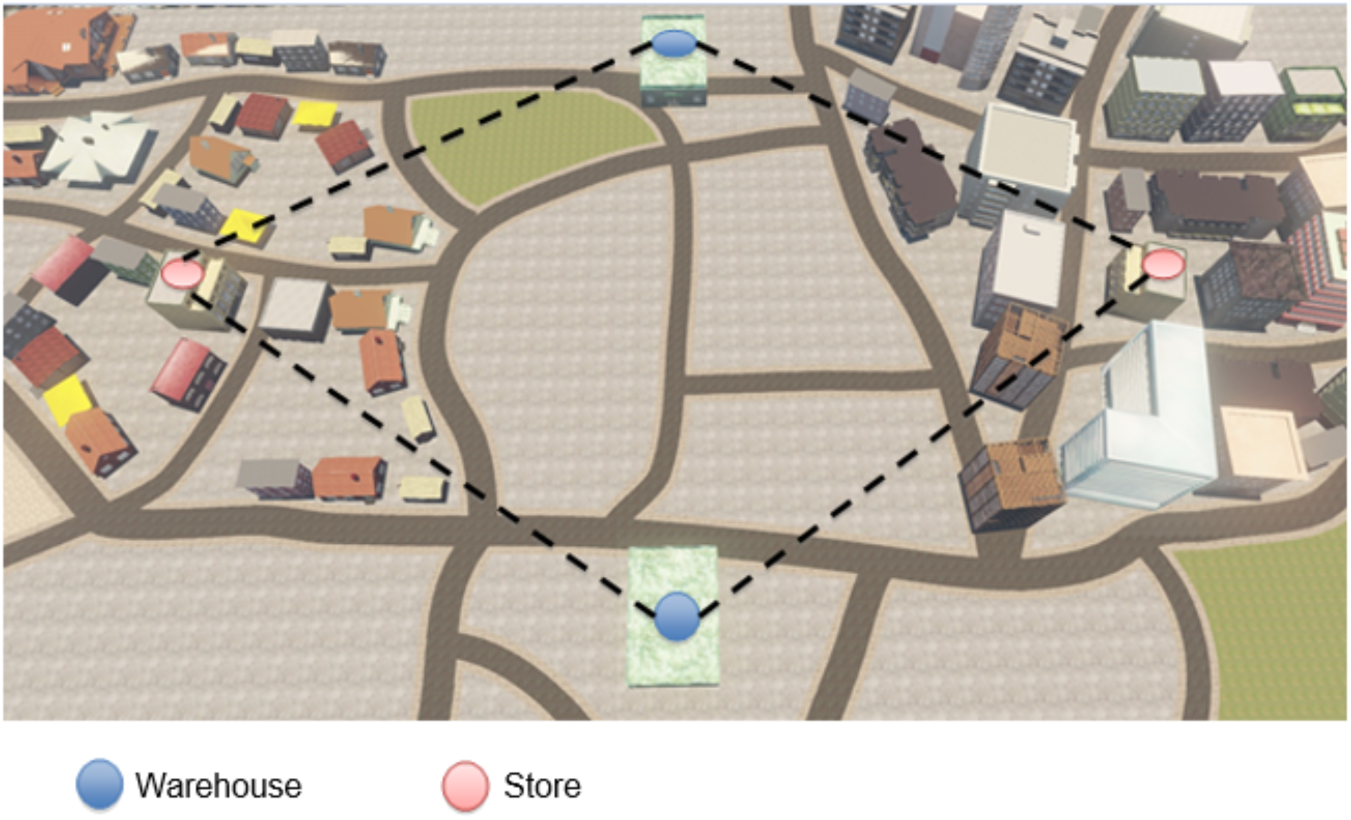
Layout of the virtual city in the spatial task. The virtual city consisted of two neighborhoods, one depicting a city center mostly with skyscrapers (right) and the other one depicting a residential area mostly with one- and two-story houses (left). The neighborhoods were separated by fallow land. On the fallow land, two identical warehouses (blue dots) were situated opposite each other. Participants had to pick up objects from one of the two warehouses and deliver it to the target store (red dots) of the corresponding neighborhood. The stores looked identical. Furthermore, Euclidean distances between the warehouses and the stores were identical (dashed black lines). The virtual city was created using Unreal Development Kit 3 (Unreal Engine 3, Epic Games, Inc.).

**Supplementary Fig. 3.**
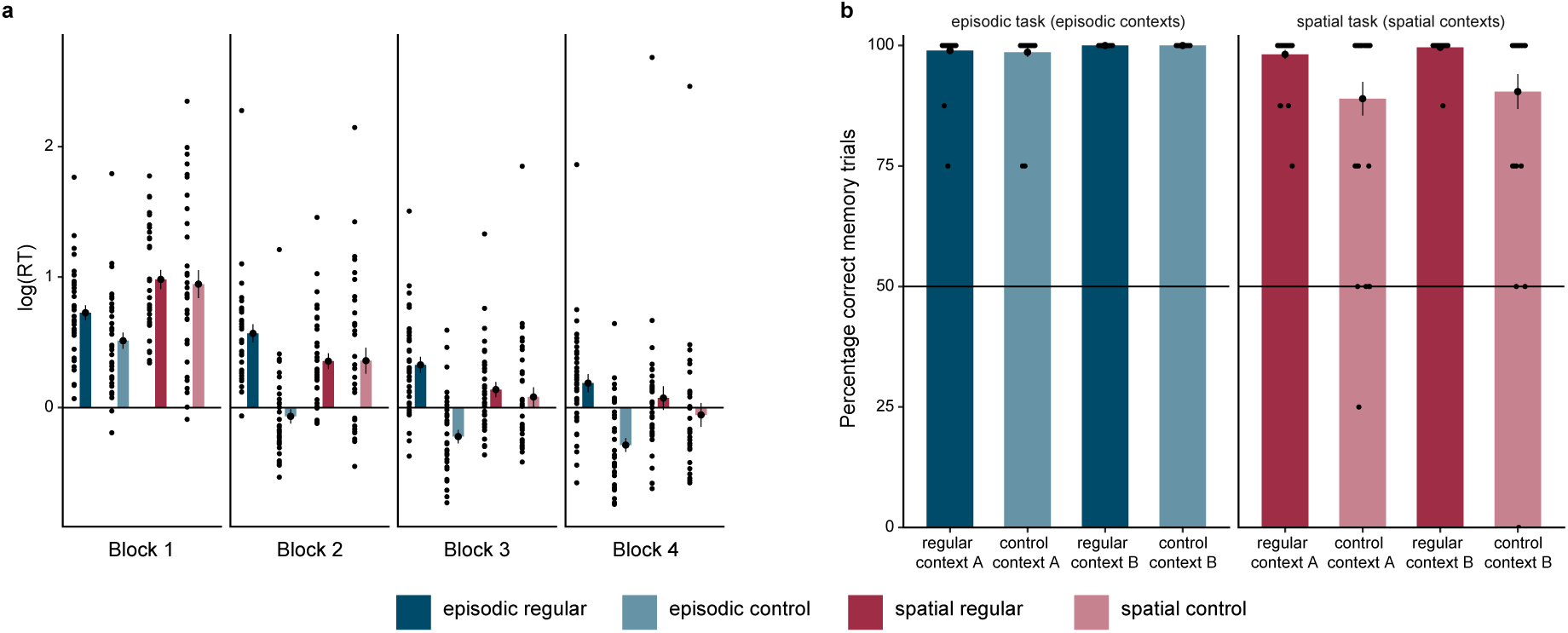
Participants form strong episodic and spatial context associations. **a** Reaction times in the memory tests after each task block (log-transformed). Reaction times are depicted separately for the episodic and spatial task and for regular and control objects. There was a significant interaction of task x block for regular objects (*F*(3,99) = 13.25, *p* < .001) but not for control objects (*F*(3,99) = 1.98, *p* = .12; main effect block: *F*(3,99) = 114.90, *p* < .001; main effect task: *F*(1,33) = 21.93, *p* < .001). **b** Performance in the final memory tests, depicted separately for the four contexts (two episodic and two spatial contexts) and for regular and control objects. For regular objects, there was no significant difference in performance between the four contexts (*F*(3,99) = 1.72, *p* = .17). For control objects, there was a significant difference in performance between the four contexts (*F*(3,99) = 6.36, *p* < .001), driven by lower performance for spatial control objects. **a,b** Dots represent data from *n* = 34 participants; bars and black circles with error bars correspond to mean ± SEM.

**Supplementary Fig. 4.**
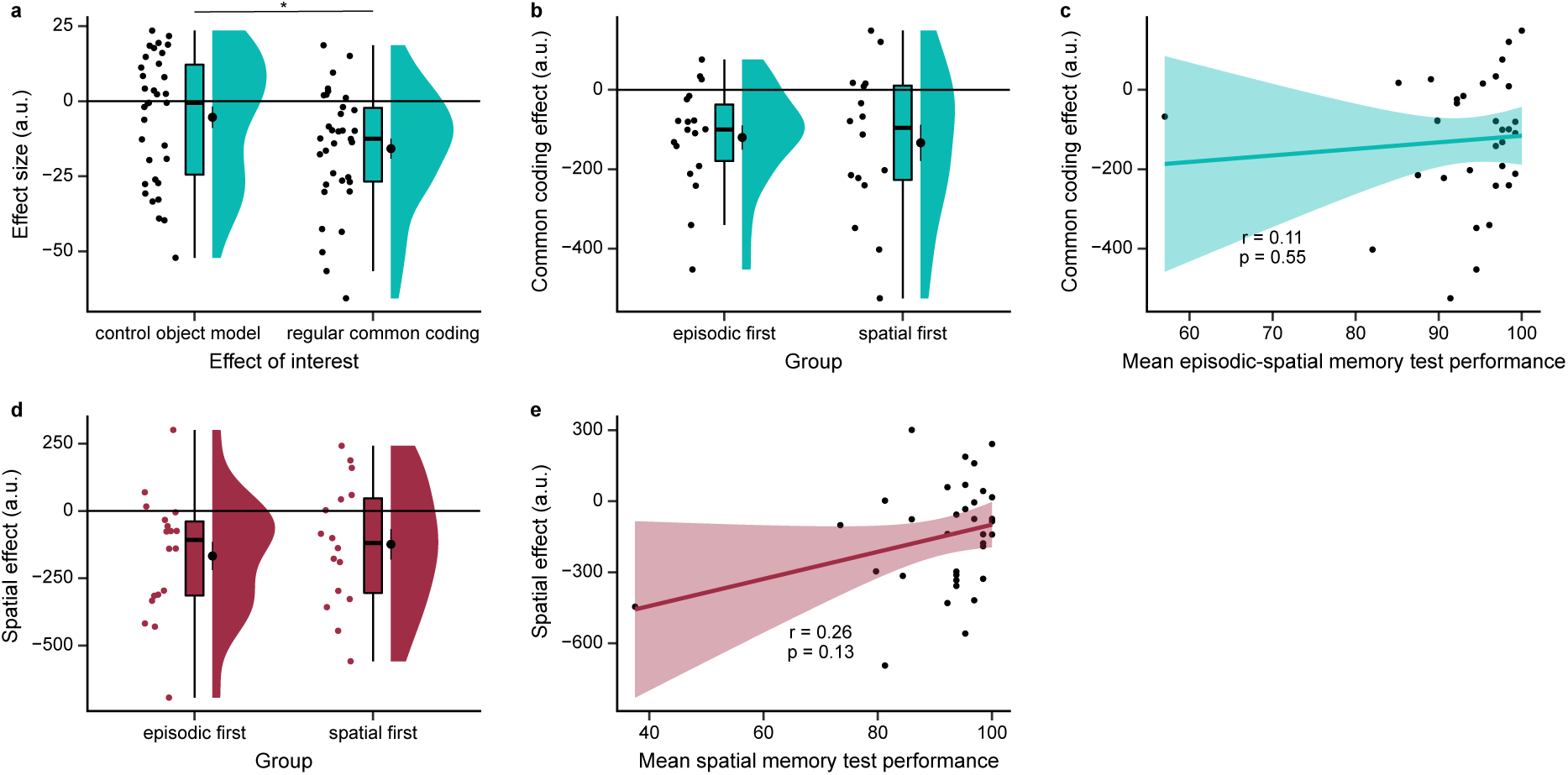
Common coding and parallel processing mechanism for episodic and spatial memory in the hippocampus. **a** Control analysis comparing the common coding effect in the anterior hippocampus with the adaptation effect for control objects. Since the episodic and spatial context relationships between objects in the common coding model scaled with the number of context associations (no, one and two), we controlled for an effect of association strength using control objects. Control objects either share two context associations in the same task (episode or space) or no context association. However, we found no significant adaptation effect for control objects (*t*(33) = -1.48, *p* = .08; one-sided test) in the common coding cluster. Furthermore, the common coding effect for regular objects was significantly stronger than the effect for control objects (*t*(33) = -2.26, *p* = .03). For comparability of the two conditions, the effect sizes of the relevant parametric regressors were averaged for each condition. **b** The common coding effect in the anterior hippocampus did not differ by group of participants having completed the episodic or spatial task first (*t*(32) = 0.24, *p* = .80). **c** There was no significant correlation between the common coding effect and performance in the episodic and spatial memory tests, averaged across tasks and blocks (note that performance was at ceiling in the final test; also n.s. without outlier). **d** The spatial effect in the right hippocampus did not differ by group of participants having completed the episodic or spatial task first (*t*(32) = -0.55, *p* = .59). **e** There was no significant correlation between the spatial effect and performance in the spatial memory tests (averaged across blocks; also n.s. without outlier). **a,b,d** Dots represent individual participants’ data; boxplots show median and upper/lower quartile with whiskers extending to the most extreme data point within 1.5 interquartile ranges above/below the quartiles; black circles with error bars correspond to mean ± SEM; distributions depict probability density functions of data points. **c,e** Dots represent data from *n* = 34 participants; line represents linear regression line, with shaded regions as the 95% confidence interval.

**Supplementary Fig. 5.**
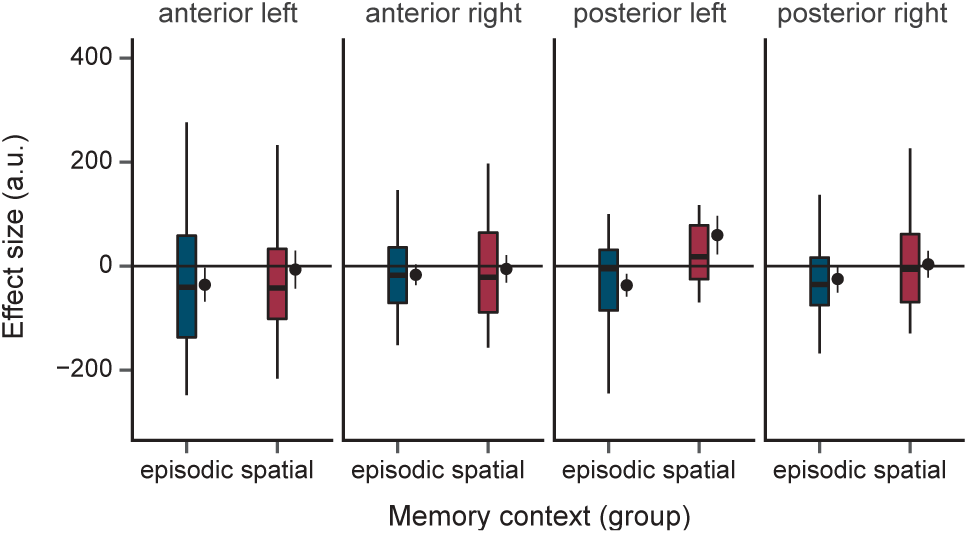
Episodic and spatial effects in PVT1. We also tested our prediction of a parallel processing mechanism in PVT1 after participants completed only one task (18 participants for the episodic task, 16 participants for the spatial task). We found no significant difference between episodic and spatial effects nor pure episodic or spatial effects (hippocampal small volume analysis, non-parametric permutation test with TFCE: difference: *pFWE* = .44; episodic: *pFWE* = .13, spatial: *pFWE* = .66, episodic and spatial one-sided tests). In addition, we conducted a complementary ROI analysis with hippocampal subregions to test for differences between episodic and spatial effects with respect to the hemispheres and the longitudinal axis (depicted in this figure). We observed no significant interactions between episodic and spatial effects with the hemisphere or longitudinal axis (Fig. 3f; interaction group x hemisphere x axis: *F*(1,32) = 1.45, *p* = .24; interaction group x hemisphere: *F*(1,32) = 2.37, *p* = .13; interaction group x axis: *F*(1,32) = 1.49, *p* = .23; interaction hemisphere x axis: *F*(1,32) = 2.38, *p* = .13; group: *F*(1,32) = 1.50, *p* = .23; hemisphere: *F*(1,32) = 0.18, *p* = .67; axis: *F*(1,32) = 0.92, *p* = .35). Boxplots show median and upper/lower quartile with whiskers extending to the most extreme data point within 1.5 interquartile ranges above/below the quartiles; black circles with error bars correspond to mean ± SEM.

**Supplementary Fig. 6.**
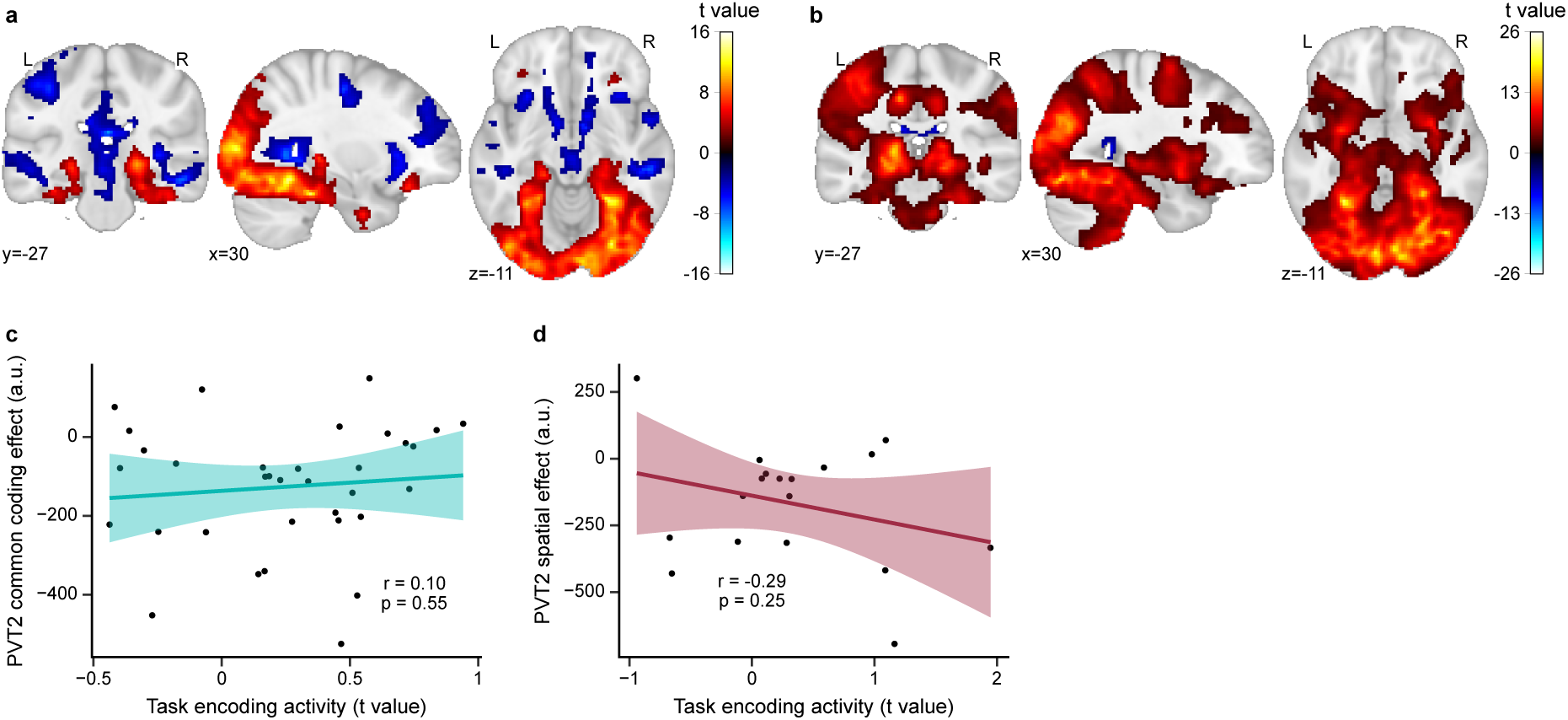
Encoding-related activity during the context association tasks. Participants learned the object-context associations during the episodic life-simulation task and the spatial virtual reality task. They performed one of these tasks in the fMRI scanner (16 participants for the episodic task and 18 participants for the spatial task). We explored encoding-related activity during these tasks and tested whether this would correlate with the adaptation effects in PVT2. **a** Encoding-related activity in the episodic task while participants watched the video of the fictional character performing the object-associated action. **b** Encoding-related activity in the spatial task while participants delivered the object to the neighborhood. **c** There was no significant correlation between encoding-related activity in the episodic and spatial tasks and the common coding effect in PVT2 in the anterior hippocampus. **d** There was no significant correlation between encoding-related activity in the spatial task and the spatial effect in PVT2 in the right hippocampus. **a-b** Non-parametric permutation tests with TFCE and *pFWE* < .05. Statistical images are displayed on the MNI template. **c-d** Dots represent data from *n* = 34 (c) and *n* = 18 (d) participants; line represents linear regression line, with shaded regions as the 95% confidence interval. Note that we used *t*-values for the episodic and spatial task effects for the purpose of task comparability.

## Acknowledgements

We would like to thank Lorena Deuker for her valuable input on the study design and setting up technical scripts for the study. We want to thank colleagues of the Doellerlab for helpful discussions of the study. This work was funded by a NWO Top Talent grant (406-15-291) awarded to N.d.H. and by the Max Planck Society. CFD is supported by the Max Planck Society, the European Research Council (ERC-CoG GEOCOG 724836), the Kavli Foundation, the Jebsen Foundation, Helse Midt Norge and The Research Council of Norway (223262/F50, 197467/F50).

## Data and code availability

Data and code to reproduce the figures and statistical analyses will be made openly available.

## Author contributions

A.N. and N.d.H. contributed equally to this work. The following list of author contributions is based on the CRediT taxonomy. Conceptualization: N.d.H., C.F.D.; Data curation: N.d.H., A.N.; Formal analysis: A.N., N.d.H.; Funding acquisition: N.d.H., C.F.D.; Investigation: N.d.H., A.N.; Methodology: N.d.H., A.N., M.M.G, C.F.D.; Project administration: N.d.H., A.N., C.F.D.; Resources: N.d.H., A.N., C.F.D.; Software: N.d.H., A.N.; Supervision: C.F.D., M.M.G.; Validation: A.N., N.d.H.; Visualization: A.N., N.d.H.; Writing -original draft: A.N., N.d.H.; Writing - review & editing: A.N., N.d.H., M.M.G., C.F.D.

